# Behavioral diversity assisting obstacle navigation during group transportation in ant

**DOI:** 10.1101/194100

**Authors:** Shohei Utsunomiya, Atsuko Takamatsu

**Affiliations:** Department of Electrical Engineering and Bioscience, Waseda University, Shinjuku-ku, Tokyo 169-8555, Japan

## Abstract

Cooperative transportation behavior in ants has attracted attention from a wide range of researchers, from behavioral biologists to roboticists. Ants can accomplish complex tasks as a group whereas individual ants are not intelligent (in the context of thisstudy’s tasks). In this study, group transportation and obstacle navigation in *Formica japonica*, an ant species exhibiting ‘uncoordinated transportation’ (primitive group transportation), are observed using two differently conditioned colonies. Analyses focus on the effect of group size on two key quantities: transportation speed and obstacle navigation period. Additionally, this study examines how these relationships differ between colonies. The tendencies in transportation speed differ between colonies whereas the obstacle navigation period is consistently reduced irrespective of the colony. To explain this seemingly inconsistent result in transportation speed, we focus on behavioral diversity in ‘directivity’, defined as the tendency of individual ants to transport a food item toward their own preferential direction. Directivity is not always toward the nest, but rather is distributed around it. The diversity of the first colony is less than that of the second colony. Based on the above results, a mechanical model is constructed. Using the translational and rotational motion equations of a rigid rod, the model mimics a food item being pulled by single or multiple ants. The directions of pulling forces exerted by individual ants are assumed to be distributed around the direction pointing toward the nest. The simulation results suggest that, as diversity in directivity increases, so does the success rate in more complicated obstacle navigation. In contrast, depending on group size, the speed of group transportation increases in the case of lower diversity while it is almost constant in the case of higher diversity. Transportation speed and obstacle navigation success rate are in a trade-off relationship.

## Author summary

Cooperative transportation behavior in ants attracts researchers from behavioral biologists to roboticists. Ants can accomplish complex tasks in groups, whereas individuals are not likely intelligent. Recent reports focus on higher-level mechanisms in coordinated transportation, e.g., triggering by an informed ant. In contrast, thisstudy focuses on the simplest form of group transportation where there is no specialist in the group. We analyze group transportation and obstacle navigation combined with mechanical model analysis. Behavioral diversity in ‘directivity’ is found to govern performance effectiveness. Here, directivity is defined as the tendency of an individualant to preferentially transport a food item in its own direction rather than in the nest direction. Such outliers can impede transportation speed but contribute well to the system when a group of ants is hindered by a complex obstacle. It is noteworthy that the two measures of efficiency, transportation speed and success rate of obstacle navigation, are in a trade-off relationship.

## Introduction

Fully coordinated systems are fragile because, once a part of the system becomes unstable, the whole system can sometimes cease to function. In contrast, a fully decentralized system consisting of simple elements can work well even if some elements are broken. Group transportation in ants is a good example of such a decentralized system. Although individual ants are not intelligent in the context of this study ‘s tasks, they can accomplish complex tasks through local interaction, sometimes even without a leader. Consequently, researchers from fields as diverse as behavioral biology, physics, and robotics have studied group transportation in ants [1–7]. In contrast, quantitative analysis of group transportation behavior has been insufficient until recently.

Group transportation is a behavior in which two or more worker ants move a large food item that not transportable by a single ant. The behavior was categorized into three kinds or transportation by Czaczkes and Ratnieks: uncoordinated, encircling coordinated, and forward-facing cooperative [1]. In uncoordinated transportation, ants pull a food item in opposite directions [8], resulting in slow-speed transportation and/or frequently reaching a deadlock. In encircling coordinated transportation, ants encircle a food item and the majority of members align at the side of the moving front and pull it, whereas minor members align at the back and pull, lift, or push it. This behavior achieves quick transportation, rarely reaching a deadlock. In forward-facing cooperative transportation, a leader ant lifts the food item as its head faces forward while smaller ants subsequently join behind the leader to stabilize the transportation of the item [9]. In the last case, the role and appearance of each ant are clearly distinct; thus, it is out of scope in this study. Uncoordinated transportation has been categorized by McCreery and Breed [10] as transportation without ‘information transfer,’ and the latter two as transportation with ‘information transfer.’

Recently, encircling coordinated transportation or transportation with information transfer has been investigated intensively in cases where worker ants appear indistinct but have roles that seem to be different during transportation. The workers of the longhorn crazy ant, *Paratrechina longicornis*, exhibit fluctuating motion during group transportation. However, they switch to straight movement when closer to the nest once a single informed ant, possessing information on the direction of the nest, attaches itself to the group [3]. Furthermore, this species exhibits excellent performance during obstacle navigation, switching its behavior intrinsically without the attachment of an informed ant [11]. Moreover, under imposition of a constraint such as an obstacle barrier, oscillatory group motion emerges which assists in navigating the obstacle [12].

To achieve the above higher-level strategies for group transportation and obstacle navigation, sophisticated mechanisms are required. In contrast, this study focuses on a lower-level but very simple strategy. By observing the behavior of Japanese wood ants, *Formica japonica*, the fundamental problem emerging through local interaction was investigated, where individual worker ants follow a simple rule and interact through the food item according to kinematics.

The ants, *F. japonica* are widely distributed in Japan [13, 14] and have been investigated particularly with regard to landmark navigation [15], social interaction [16, 17], etc. Most ants of this species lives under open ground and transport large food items in groups without dissection [18]. During group transportation, the ants pull the food item in their own preferential directions (Fig 1);Some pull the item toward their nest (denoted by a black arrowhead) and others pull it in the opposite direction of the nest (denoted by a white arrowhead). This results in tug-of-war movement (S1 Video). Thus, their manner of transportation could be categorized as ‘uncoordinated’ [1] and this species has been believed to exhibit low-efficiency transportation [1, 8, 10]. This is, however, a good model animal for investigating how the lowest-level strategy can solve tasks using simple rules. We show that the key mechanism is behavioral diversity in ‘directivity, ‘which is defined as the tendency of an individual ant to transport a food item in its own preferential direction. How diversity in directivity supports obstacle navigation is discussed.

**Fig 1.**
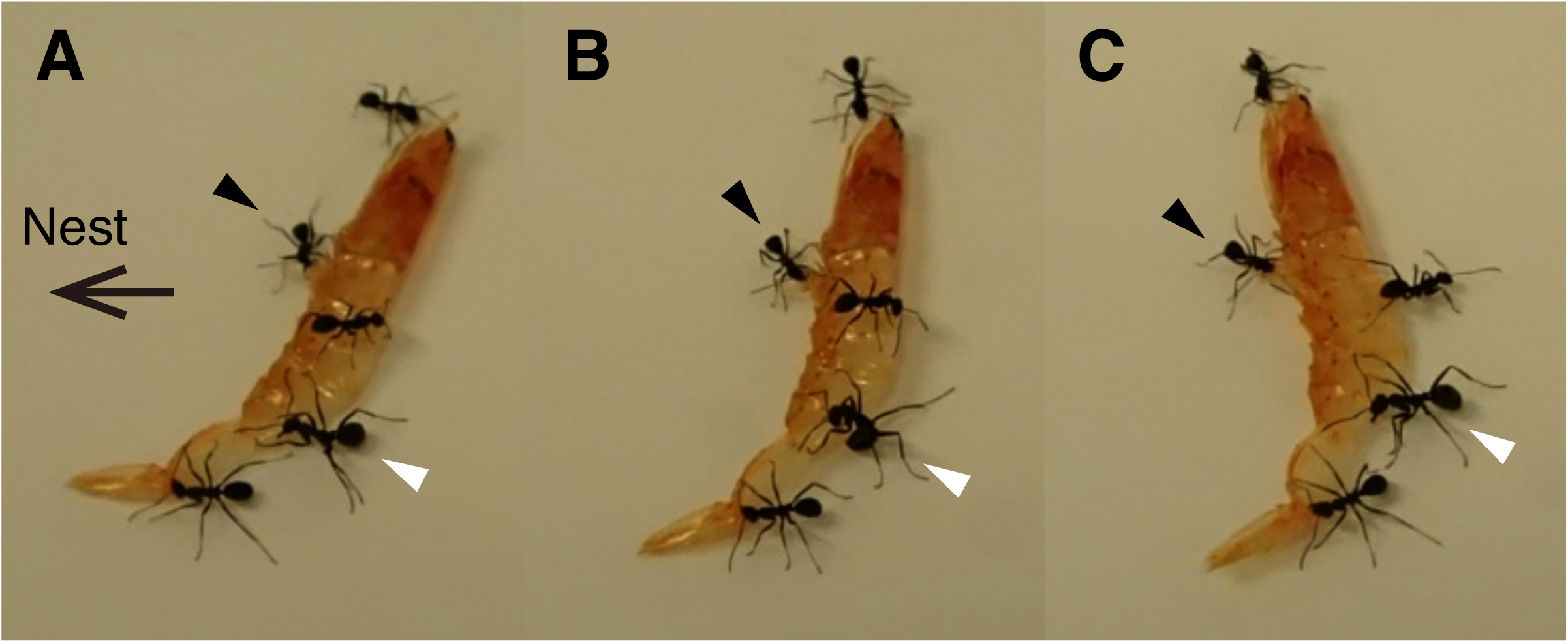
Tug of war observed in ants, *Formica japonica*, transporting a food item. (A–C) are arranged chronologically in 1-s intervals. The arrow indicates the direction of the nest. The ant denoted by the black arrowheads attempts to drag the food item toward the nest while the ant denoted by the white arrowheads acts against it, resulting in moving the food item in the opposite direction from the nest. See also S1 Video.

## Results

The behavior of *F. japonica* ants during group transportation of a food item toward their nest and their obstacle navigation were observed on a stage placed in an indoor artificial environment termed colony (I), or in the outdoor natural environment termed colony (II) (Fig 2; See the Materials and Method section for details). Then the ransportation speed, obstacle-passing period, and direction of transportation were analyzed.

**Fig 2.**
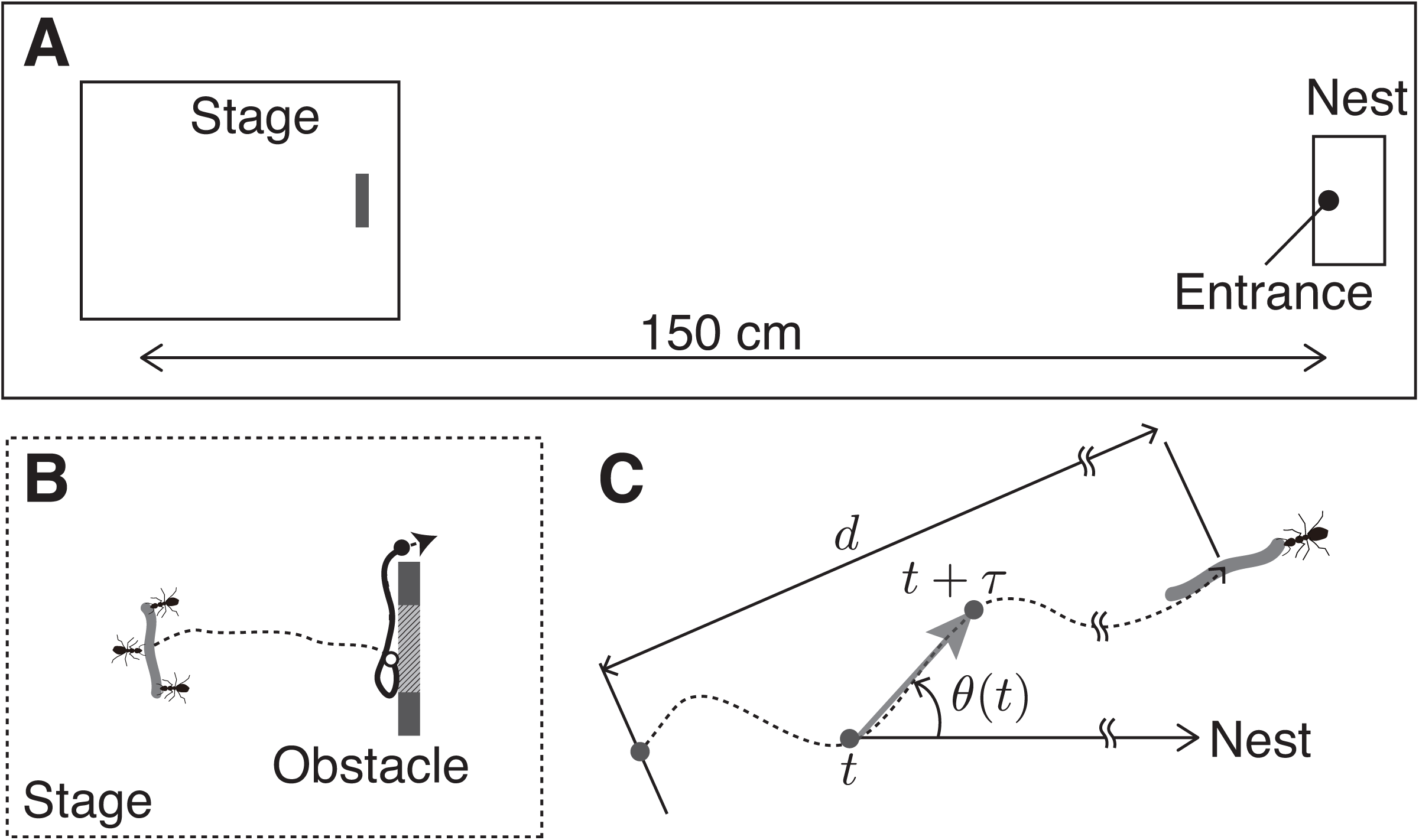
Experimental setup. (A) Setup for observation. An observation stage (W40 × D 30 cm) equipped with a ‘rat-proof ‘structure was placed 150–200 cm away from the nest entrance. (B) An obstacle made of polystyrene (W 6.8 × D 1.5 × H 3.7 cm) was placed in front of the ants transporting a food item. Open and closed circles represent positions where an ant group touched the obstacle and passed through it,respectively. The part shaded with oblique lines on the obstacle represents the target area that one of the ants touch it first, whose length was 3.4 cm. (C) Illustration for data analysis.

In individual transportation, ants moved the food items, which were 5–10 times longer and 20 times heavier than a worker ant, by pulling them. During long periods of group transportation with two or more members, the composition of the members was dynamically substituted as reported in the above literature [3, 11, 19]. This behavior can cause randomness, fluctuating behavior, and/or ‘information transfer’ into the group by newly attached ants. In contrast, in this study only cases of fixed member composition were analyzed because we focused on the basic level of group behavior.

### Transportation in a group and obstacle navigation

The transportation speed of a food item by multiple ants in a group depended on the group size in colony (I), as shown in Fig 3A, while it did not in colony (II), as shown in Fig 3B. When the ant group encountered an obstacle placed on the way to the nest, they initially struggled but finally succeeded in passing through the obstacle(S2 Video). The obstacle-passing period showed a decreasing tendency depending on the group size in both colonies (Fig 3C and D), although the tendency is unclear in the results of colony (II). It is noteworthy that 25% (colony (I); denoted as “Abn.” in Fig 3E) and 67% (colony (II)) of transportation trials by single ants failed; that is, the ants finally abandoned the task and released their food items (S3 Video), whereas the groups consisting of more than two ants rarely abandoned their food items (Fig 3E). The distribution of the obstacle-passing period by single ants was broad (Fig 3C). This could be caused by the variation of pulling power among individual ants. The passing period in transport by an individual ant decreased depending on speed and ranged from 5 to 45 s, whereas in multiple-ant group transportation, it reduced to less than 25 s, and there was no significant dependency on transportation speed (Fig 3F). Taken together, it can be concluded that transportation by more than two ants reduces the period passing the obstacle and increases the probability of success.

**Fig 3.**
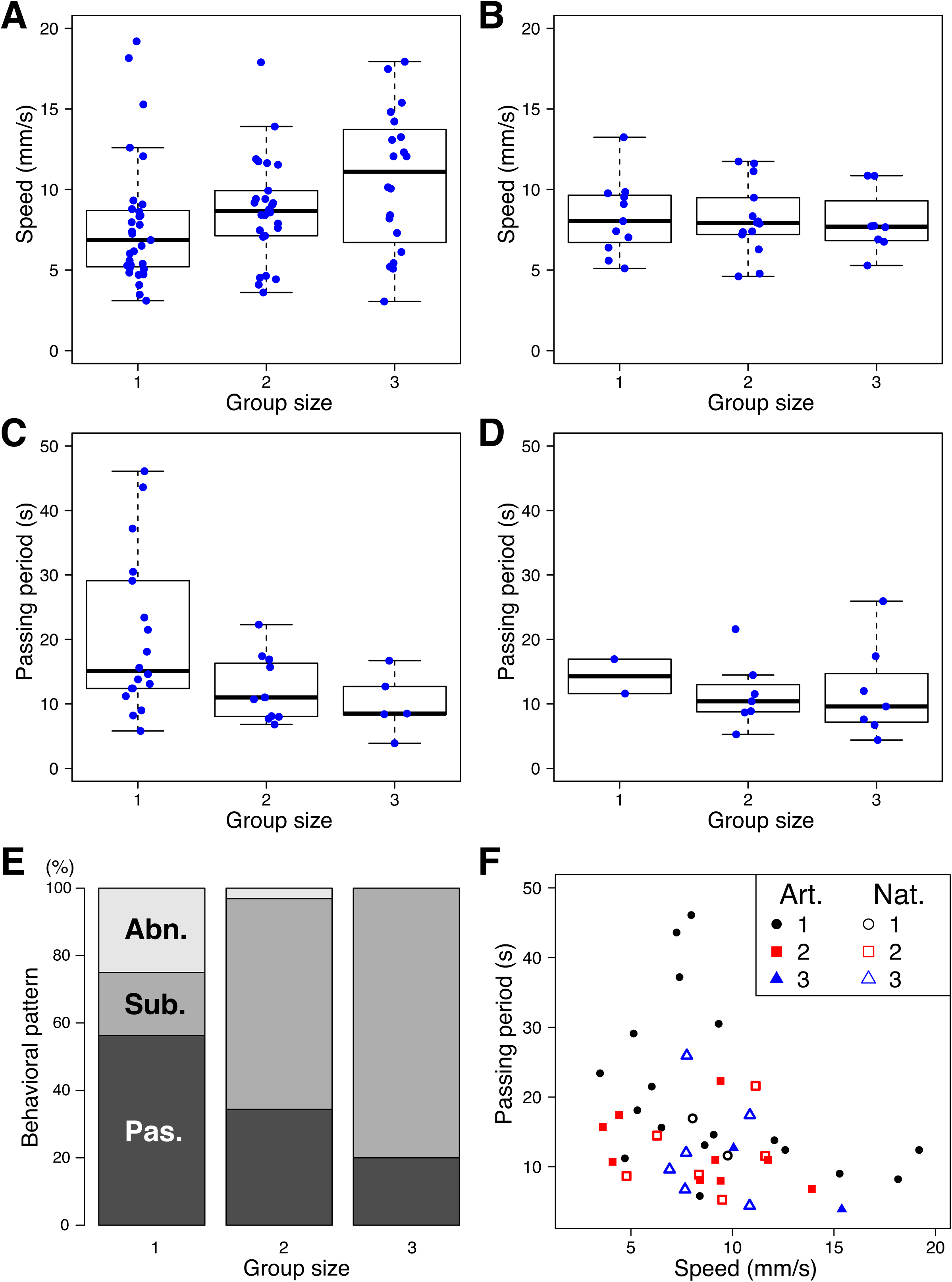
Transportation in a group and obstacle navigation. (A and B) Dependence of transportation speed on group size. (C and D) Dependence of obstacle-passing period on group size. (E) Behavioral patterns when a group of ants faces an obstacle. “Pas.” denotes the case where the group succeeded in the task with its initial members and arrangement. “Sub.” denotes the case where at least one of the members left and/or was substituted for another ant but where the group eventually succeeded in the task. “Abn.” denotes the case where the members abandoned the transportation task and released the food item. Only the “Pas.” cases are included in C and D. Trials of A, C, and E were performed in the indoor artificial environment (colony (I)). Trials of B and D were performed in the outdoor natural environment (colony (II)). (F) Relation between transportation speed and obstacle-passing period. Closed and open marks represent data obtained from colonies(I) and (II), respectively. Black circles, red squares, and blue triangles denote group sizes of 1–3, respectively. In box-and-whisker diagrams, the bottom and top of the boxes denote the first and third quartiles, respectively; the thick band inside the box shows the median; the end and the top of the whiskers represent the lowest and highest data points within 1.5 times the interquartile range (IQR). Scatter plots show the original data. All data sets in A and B passed normality and equal variance tests. Part of the data sets in C and D did not pass them. Thus, both one-way ANOVA and Kruskal-Wallis (KW) tests were applied. A: ANOVA, *p* = 0.04838 *<* 0.05; KW,*p* = 0.03399 *<* 0.05. B: ANOVA, *p* = 0.957 *>* 0.05; KW, *p* = 0.9102 *>* 0.05. C:ANOVA, *p* = 0.004097 *<* 0.05; KW, *p* = 0.05354 *<* 0.1. D: Statistical analysis was not applied because the size of the data for a group size of 1 is too small (*n* = 2).

### Directivity during transportation

To investigate the behavioral variation of individual ants, directions of food item transported single ants were analyzed. Fig 4 A shows an example of a trajectory of a food item transported by a single ant. The ant transported it in a certain direction while keeping an almost constant angle relative to the nest direction. The direction angle was defined as the directivity of the ant using *θ*_0_. It was estimated using the mean value of the transportation directions *θ* at each measurement time, represented as a histogram in Fig 4B. In the mean value calculation, the outliers were excluded; these correspond to sudden changes of direction denoted by black arrow heads in Figs 4A and B (see also the Materials and Method section). The cause of these sudden changes is unknown; however, they could occur when part of the food item is caught by friction with the ground, or by some internal mechanism of the ant. Even after excluding these exceptions, the transportation direction still fluctuated around *θ*_0_,which was defined as the magnitude of fluctuation *σ* equal to the standard deviation of the *θ* distribution (assumed to be normal distribution and represented with a red line in Fig 4B). The median values of the *σ* distribution were 12.1 (10.24, 15.6) and 13.5(7.7, 19.9), in colonies (I) and (II), respectively, where the first and second values in parentheses represent the first and third quartiles of the distribution, respectively.

**Fig 4.**
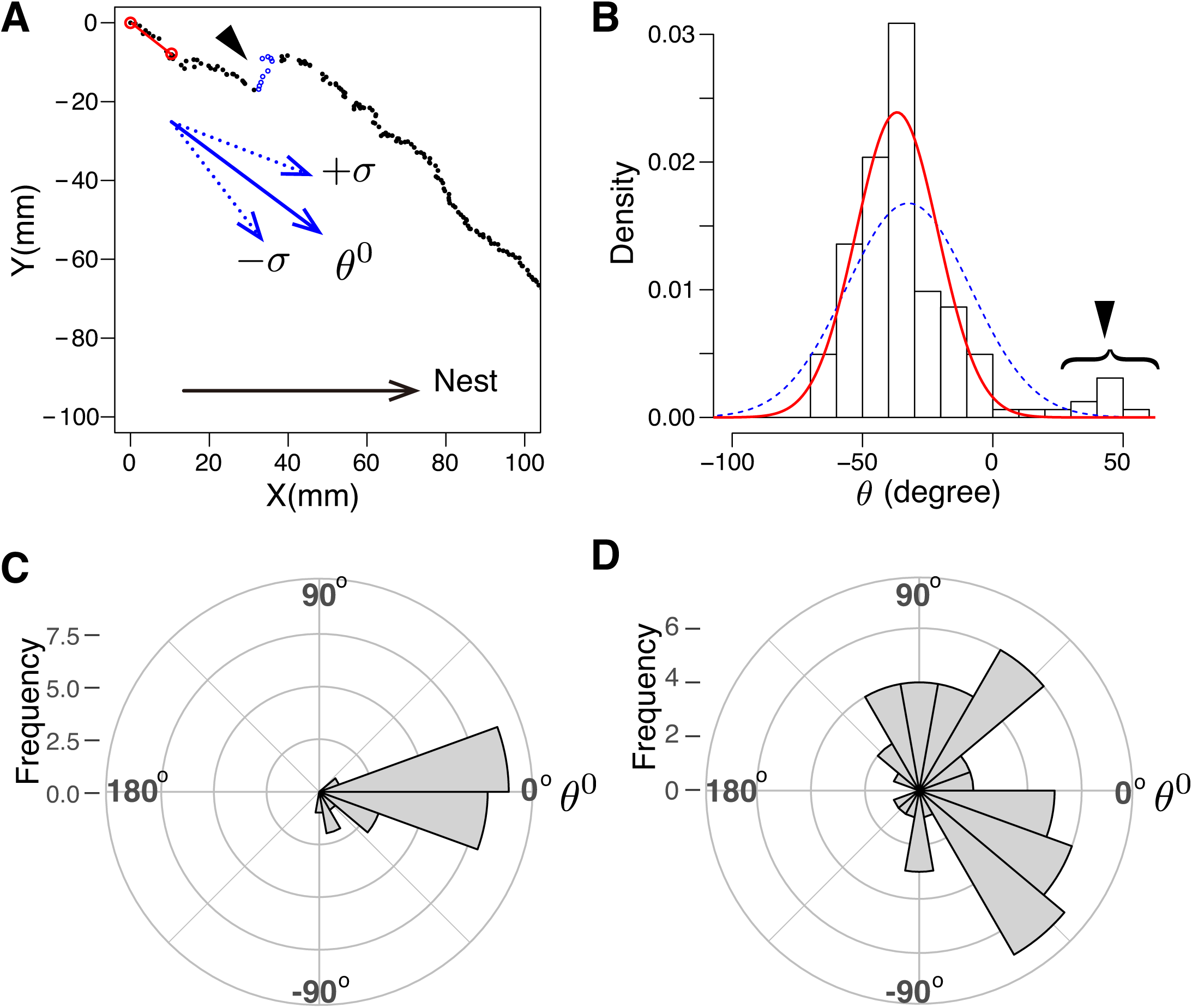
Directivity *θ*_0_ and fluctuation *σ* in trajectories during transportation by single ants. (A) An example of trajectory of the food item transported by a single ant. The time intervals of the data denoted by black closed circles are *δt* = 0.1s. The time duration for estimation of the transportation direction is *τ* = 1.0 s, whose example is denoted by red open circles and a red line (see also the Materials and Methods section and Fig 2C). (B) Distribution of the transportation directions *θ*relative to the direction of the nest, estimated using the same data shown in Fig 4A. The distribution was fitted with a normal distribution using all the data (blue dashed line) and using partial data excluding outliers (red solid line; see also the Materials and Methods section). The mean value of the latter data was defined as the directivity *θ*_0_ of an ant and the standard deviation as the magnitude of fluctuation *σ*. The data in Fig 4A and B were obtained from colony (I). (C and D) Directivity *θ*_0_. Data shown in Figs 4C and D were obtained from colonies (I) and (II), whose population sizeswere *n* = 25 and *n* = 50, respectively.

The directivity *θ*_0_ was quite persistent for individual ants (S4 Video). Thus, *θ*_0_ can determine individuality for each ant. Figures 4C and D show the distributions of *θ*_0_ in colonies of (I) and (II), respectively, in which *θ*_0_ scatters around the direction of thenest (*θ*_0_ = 0). The mean values of *θ*_0_ in colonies (I) and (II) were *-*12.48*°* and 12.27*°*, respectively. Their variances *V*, denoting the diversity of *θ*_0_, were 0.13 and 0.62 in*-*colonies (I) and (II), respectively. This quantity is defined as *V* = 1-*R* in the domain*≤ ≤*of 0≤*V*≤1 using order parameter *R* (described later in Eq (3)). *V* = 0 suggestscoherence, i.e., ants from the same colony exhibit a high degree of consensus in their moving direction; on the other hand, *V* = 1 suggests diversity, i.e., individual antsmove in dispersed directions. Surprisingly, a considerable number of ants in colony (II) transported the food item in directions quite different from that toward their nest.

## Discussion

The results of the experimental observations and analyses can be summarized as follows: (1) the speed of transportation in a group depended on the group size in colony (I) but not in colony (II); (2) the obstacle-passing period tended to decrease with group size and the success rate increased with grouping; (3) the remarkable difference in the properties of colonies (I) and (II) was in their diversity(individuality), measured by the directivity of individual ants. Thus, we hypothesize that diversity in directivity leads to effective obstacle navigation in groups and causes the different tendencies observed in transportation speed between the colonies.

To verify this hypothesis, we constructed a simple mechanical model representing ant group transportation of a food item assuming that each member possesses individuality in its directivity.

## Model

The motion of a food item in traction by single or multiple ants was modeled using translational and rotational motion equations. As already pointed out in the review by McCreery and Breed [10], interaction among ants can be achieved by mechanical force transmitted through the food item, which they referred to as ‘information transfer’ in their paper. The mechanical interaction can be expressed naturally by the mechanical model of the food item, but a mechanism for ‘information transfer’ should also be considered. One of the candidates is adaptive force generation, which is a mechanism by which the performance of individual ants increases according to the load per single ant, as reported in the literature [19]. Additionally, the force fluctuated with direction, as seen in our observations, and the motion of transportation by a single ant sometimes changed suddenly (Figs 4A and B). Note that these behaviors are related closely to ‘persistence,’ defined by McCreery and Breed as “worker’s reluctance to giving up or to change their direction of motion [10].” Therefore, we assumed that traction force exerted by a single ant is generated with fluctuation in directivity, stochasticity, and a simple adaptation mechanism as follows.

### Basic model equations

A food item transported by ants (*i* = 1, 2*, …, n*) was modeled as a rigid rod with mass *M* and moment of inertia *I* (Fig 5A). Each ant grips the rod at a position **r**_*i*_ relative to the rod ‘s mass center and drags it with a maximum traction force **f**_*i*_. We assumed adaptative force generation wherein the ants control the traction forcede pending on the speed of the food item. This is expressed by damping terms proportional to the velocity of the mass center of the rod **v** using a damping coefficient *γ*_*i*_. Then the motion of the rod is written using the following translational and rotational motion equations:

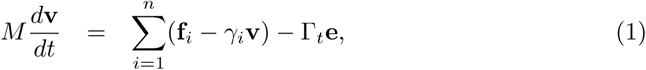

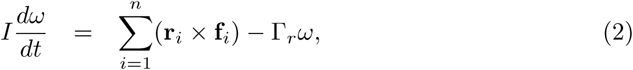

where 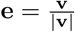 is the unit vector of the travel direction and *ω* is the angular velocity around the mass center. In both motions, kinetic friction in force and torque wereconsidered, where γ_*t*_ and γ_*r*_ are the coefficients of kinetic friction for translational and rotational movements, respectively. The latter constant was calculated as 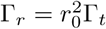 (see Eq (5) in the Materials and Methods section).

**Fig 5.**
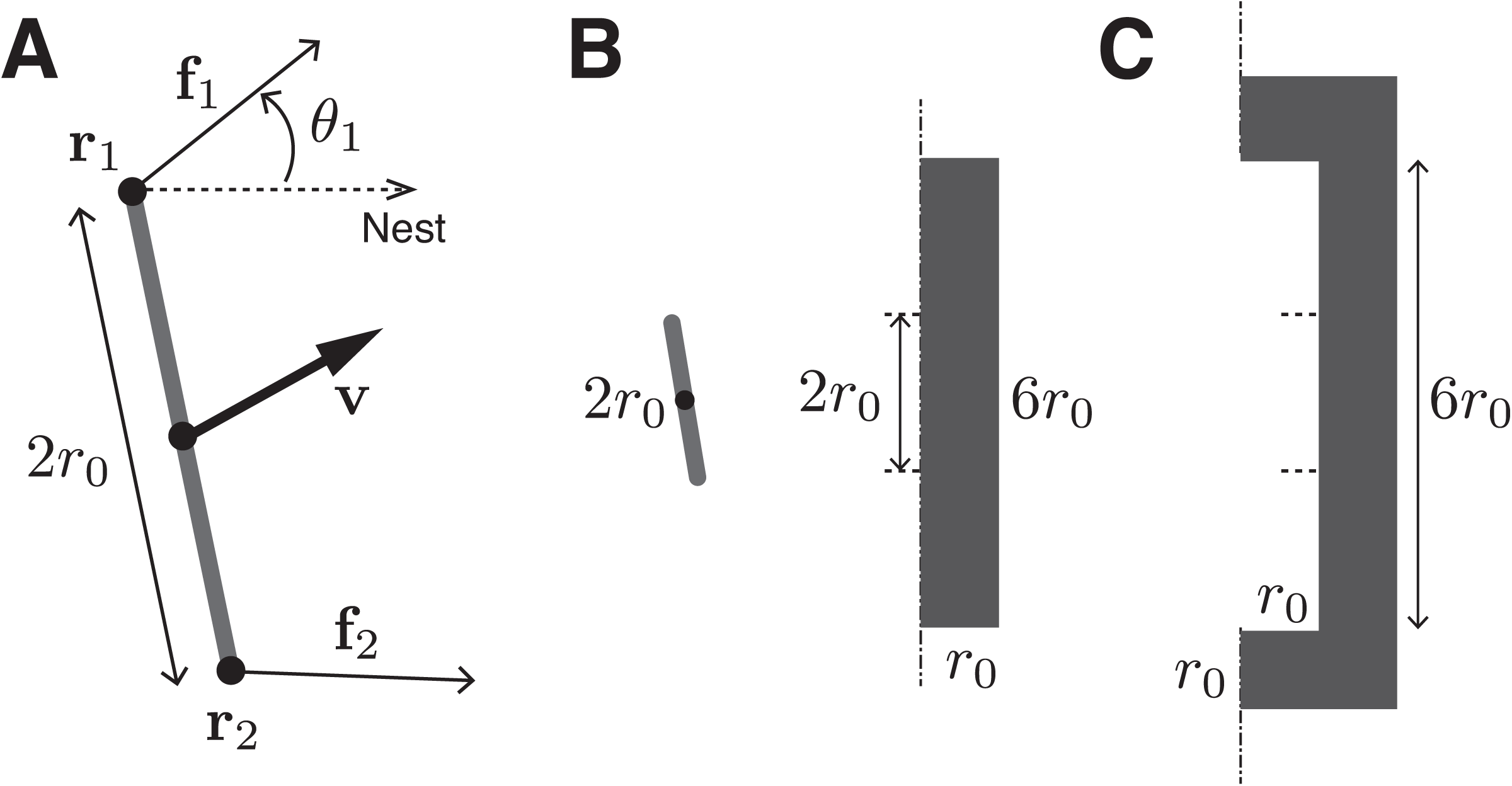
Model. (A) A rigid rod, mimicking a food item, transported by two ants. The ants exert traction forces at both ends of the rod. (B) A rectangular obstacle placed in front of the food. (C) A U-shaped obstacle. The dashdotted lines in Figs 5B and Crepresent the goal line used to judge the success of the ants in obstacle navigation.

### Directivity with fluctuating traction force

As observed in the experimental results, ants possess individual directivity when pulling a food item which is defined as 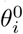. The values of 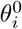 in each colony were assumed to follow a von Mises distribution, which is a circular-statistics version of the normal distribution [20] defined by two parameters: the mean and the concentration parameter *κ*. In the numerical calculations, the parameters were set with the mean equal to 0 (nest direction), and *κ* = 4.4 (for a colony with small diversity) or *κ* = 0.9(for a colony with large diversity). The parameter values were estimated roughly from the experimental results.

In addition to the individual directivity, the pulling direction fluctuated in time around 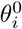 with magnitude *σ*. Therefore, the pulling direction of ant *i* at time *t* is defined as follows:

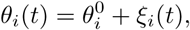

where *ξ*_*i*_(*t*) is the fluctuation of the pulling direction. The second term on the right-hand side of the equation was randomly changed in time as a Poisson process so that the mean interval was equal to *λ* = 0.1 s and the magnitude was randomly generated according to a Gaussian distribution with a mean value of 0 and a variance of *σ*^2^ = (15*°*)^2^. The variance was approximately estimated from the experimental result, and was set uniformly among the ants. Therefore, the maximum traction force is defined as follows:

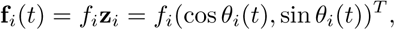

where **z**_*i*_ is the pulling direction vector.

### Ant alignment and transportation type

As reported by Czaczkes et al. [21], ants preferentially bite the corners or the protrusions of a food item. In our observations, the first and second ants tended to bite both ends of a rod-shaped food item. The others bit the remaining exposed areas. The body direction (equal to the direction of traction) of an ant’s first bite seemed sometimes to be unmatched with its preferential direction (S5 Fig). The ant pulled the food in the body-backward direction first, and later corrected the pulling direction toIts preferential direction. Buffin et al. also observed in *Novomessor cockerelli* that ants pulling the food item in the nest direction persisted to bite, but that ants biting the opposite side leave their positions more easily [19]. Based on the above observation, simulations of group transportation were performed according to the following rule.

The first and second ants bite the food item at both ends; then the other ants follow at equally spaced positions along the food item. The initial direction *θ*(0) was randomly set according to a uniform distribution, which is sometimes inconsistent with an ant ‘s preferential direction. During the simulation, ants biting at the equally spaced positions were forbidden from crossing to the opposite side of the food item.Consequently, if an ant is positioned at the side contradictory to its preferentialdirectivity *θ*_0_, the ant is not able to contribute effectively to transportation based on the above rule. In practice, such an ant would eventually leave the transportation task; then the resulting open space would become available for another ant. Thus, this case was categorized as ‘Sub.’ (for the definition, see the legend of Fig 3). Transportation speeds were estimated using all groups except for those categorized as ‘Sub.’ using the same method.

### Stochastic force generation

As seen in the Results section and in Figs 4A and B, the motion direction of transportation by a single ant suddenly changes on occasion. This could be caused by friction with the ground, resulting in stochastic force generation. The stochastic effect was introduced by setting the magnitude of the traction force *f*_*i*_ to 0 for each time step in the simulation for probability *p* = 0.25 only in the U-shaped obstacle simulation.

### Comparison with mechanical models in the literature

There have been several reports on mechanical models through physical interaction for group transportation in the literature [2, 22, 23]. In most of them, the adaptation process in individual ants was described as optimization to an assigned fitness value using an evolutionary algorithm or reinforcement learning. These approaches do not truly mimic real ant behavior but rather prioritize the effectiveness of the system as a machine.

### Model result: Speed in group transportation

In the literature, there is less consistency in the relation between speed and group size during transportation of a constant-weighted load. The speed is simply proportional to group size in *P. longicornis* [11, 12]. It is proportional at small group size and then becomes saturated at large group size in the neotropical ants *Pheidole oxyops* [21] and *Aphaenogaster cockerelli* [2]. It seems to be almost constant irrespective of group size in *Eciton burchelli* [9] and *Formica rufa* [24]. Some researches implied that larger group size causes slower speed in *Pheidole pallidula* [25] and *Novomessor cockerelli* [19] (note that the weights of the loads were not regulated in these experiments).

In contrast, two types of tendencies, proportional and constant, were found in our observations using the same species of *F. japonica* and food items of almost equal weight. This difference in tendencies could have been caused by difference in the diversity of directivity. This conjecture is verified using our model as follows.

To estimate roughly how the speed in group transportation depends on group size, Eqs (1) are rewritten by neglecting inertia as follows:

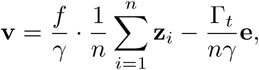

where *ξ*_*i*_(*t*) ≡ 0, *f*_*i*_ ≡ *f*, and *γ*_*i*_ ≡ γ were assumed for simplicity. The value

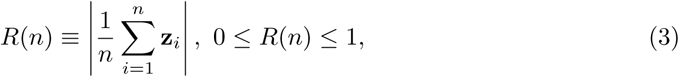

is an order parameter of directivity in the group. *R* = 0 suggests dispersal; on the other hand, *R* = 1 suggests coherence. 1 – *R* represents the degree of a group ‘s diversity *V*, as mentioned in the Results section. Because the vector **e** is the unit vector of **v**, the speed can be written as follows:

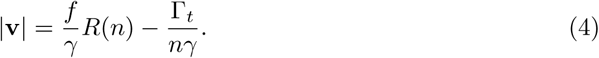

As seen in this equation, the speed consists of two converse terms: total traction force and kinetic friction force. The first term is proportional to the order parameter *R*(*n*), which depends on diversity and group size *n* as follows (see also the Materials and Methods section and S6 Fig). When there is no diversity, the order parameter is always *R*(*n*) = 1 irrespective of *n* (S6 FigA). Additionally, *R*(1) = 1 irrespective of diversity. On the contrary, when there is diversity, *R*(*n*) statistically decreases depending on *n* among a number of trials (S6 FigB and C). Larger diversity affects a larger decrease in *R*(*n*). The second term on the right-hand side of Eq (4) decreases depending on *n* irrespective of diversity. Taken together, with the above converse effects, the speed of group transportation **v** increases under small diversity, whereas it decreases or is almost constant under large diversity, as shown in the open box-and-whisker diagrams in Fig 6.

**Fig 6.**
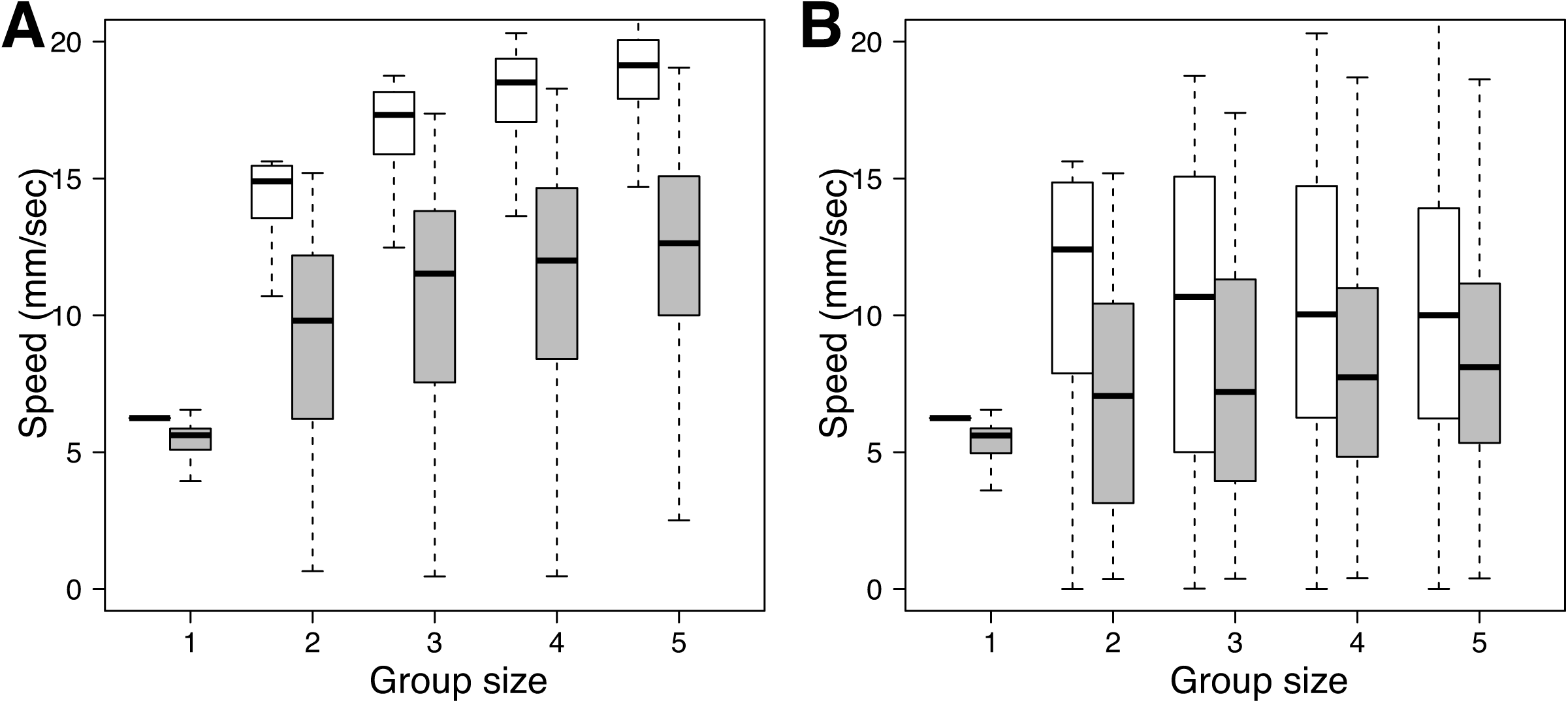
Theoretical prediction and simulation results of group-sizedependence in transportation speed. A: Small diversity in directivity (*κ* = 4.4). B: Large diversity in directivity (*κ* = 0.9). Other parameters were set as *f* = 10 gmm/s^2^, *γ* = 0.4 g/s, γ_*t*_ = 7.5 gmm/s^2^, and *r*_0_ = 10 mm. Open box-and-whisker diagrams show the results obtained using the theoretical equation Eq 4. Gray filledbox-and-whisker diagrams show the results obtained from the simulation using the mechanical model Eqs (1) and (2).

Gray box-and-whisker diagrams in Fig 6 represent the results of numerical simulation using the mechanical model of Eqs (1) and (2). The tendencies in speed dependence on group size are similar to those of the above analytical calculations;however, the absolute values were different. In fact, fluctuation of the pulling force was not considered in the theoretical equation (4), which could have caused the observed difference.

Both analyses suggest that the tendency in the speed of group transportation, whether increasing or constant, is simply governed by the diversity in directivity among individual ants belonging to each colony.

### Simulation results: passing obstacles

To investigate the relation between diversity in directivity and performance in obstacle navigation tasks by a group of ants, numerical simulations were performed (see the Materials and Methods section for details). A rod mimicking a food item being pulled by multiple ants was placed in front of a rectangular- or U-shaped obstacle, as shown in Figs 5B and C.

Figure 7 shows the result of simulation for a rectangular-shaped obstacle. When there is no diversity in directivity, the passing period was relatively longer and did not depend on group size (S7 Video, Fig 7A). When there was diversity, the passing period was reduced to almost half by grouping two ants (S8 Video, Figs 7B and C). Grouping more than two ants did not contribute further. In addition, the magnitude of diversity did not affect the results. In the simulation without diversity, 90% of single ants were not able to pass the obstacle (denoted as ‘Abn.’ in Fig 7D). The rate of ‘Abn.’ decreased by grouping two or more ants. In addition, for small diversity, the rate of ‘Abn.’ greatly decreased and was minimized for a group size two (Fig 7E). For a large diversity, the rate of ‘Abn.’ was constant irrespective of group size (Fig 7F). These findings suggest that grouping two ants with a small diversity optimize navigation of relatively simple obstacles (rectangular-shaped).

**Fig 7.**
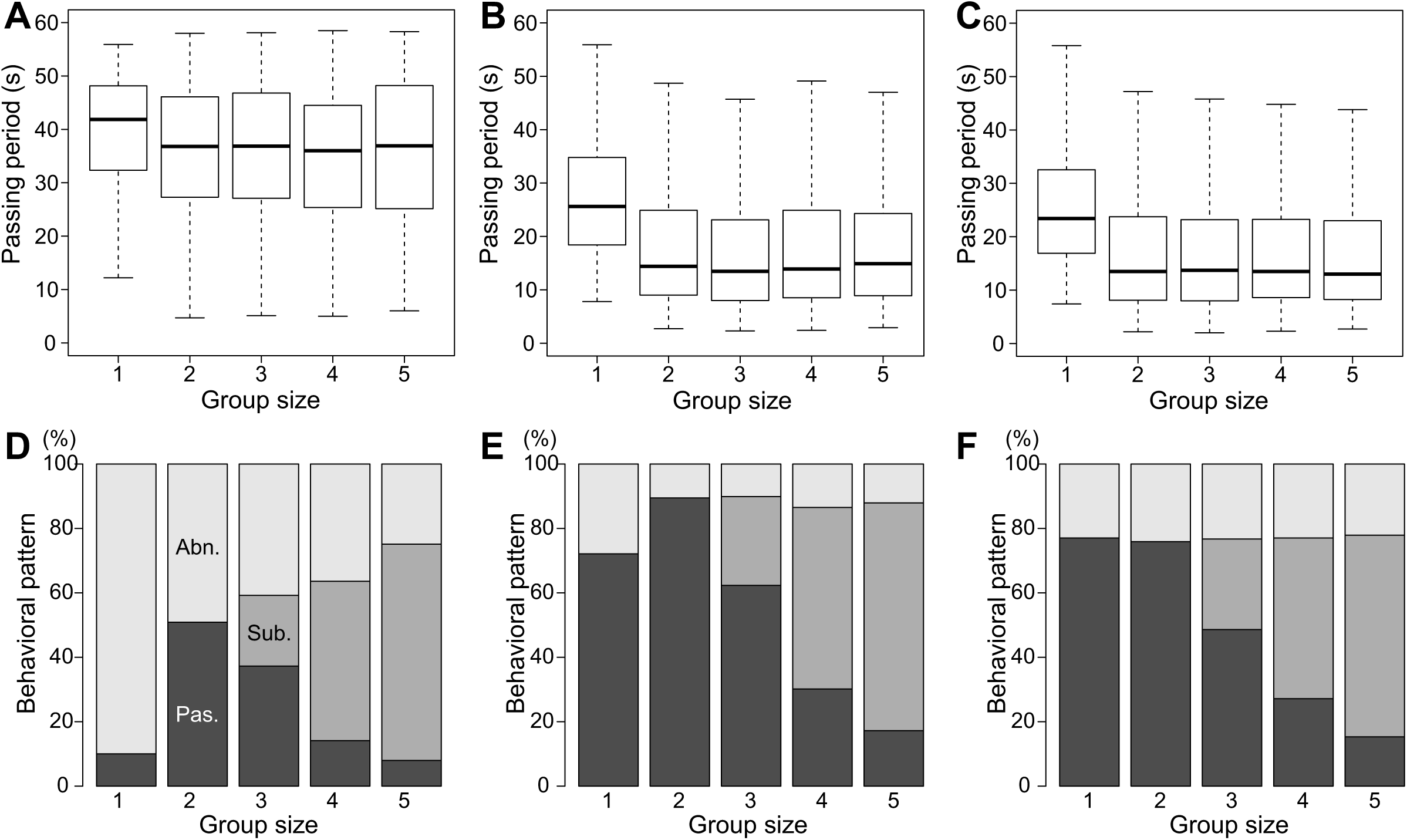
Simulation results for obstacle-passing period through a rectangular obstacle. A–C: Obstacle-passing period depending on group size. D–F: Behavioral patterns after ant groups face an obstacle. A and D: No diversity in directivity. B and E: Small diversity in directivity(*κ* = 4.4). C and F: Large diversity in directivity(*κ* = 0.9). Other parameters were set as *f* = 10, *γ* = 0.4, and γ_*t*_ = 7.5. Other notations and classification of behaviors are the same as those in Fig3.

Figure 8 shows the result of simulation for a U-shaped obstacle. In contrast to the case of the rectangular-shaped obstacle, the success rate of obstacle navigation was extremely low in the small-diversity case (‘Sub.’ and ‘Pas.’ in Fig 8D) but improved in the large-diversity case (Fig 8E). This suggests that, the larger the diversity, the more able an ant group to pass complicated obstacles. The large diversity in colony (II) in the natural environment could be effective in navigating more complex terrain. Stochastic force generation in addition to diversity in directivity moderately increased the success rate in this task (Fig 8F). In all cases, the obstacle-passing period decreased with group size except in the single-ant case. These results suggest that grouping more than two ants and the presence of large diversity values can assist in navigation of complicated obstacles.

**Fig 8.**
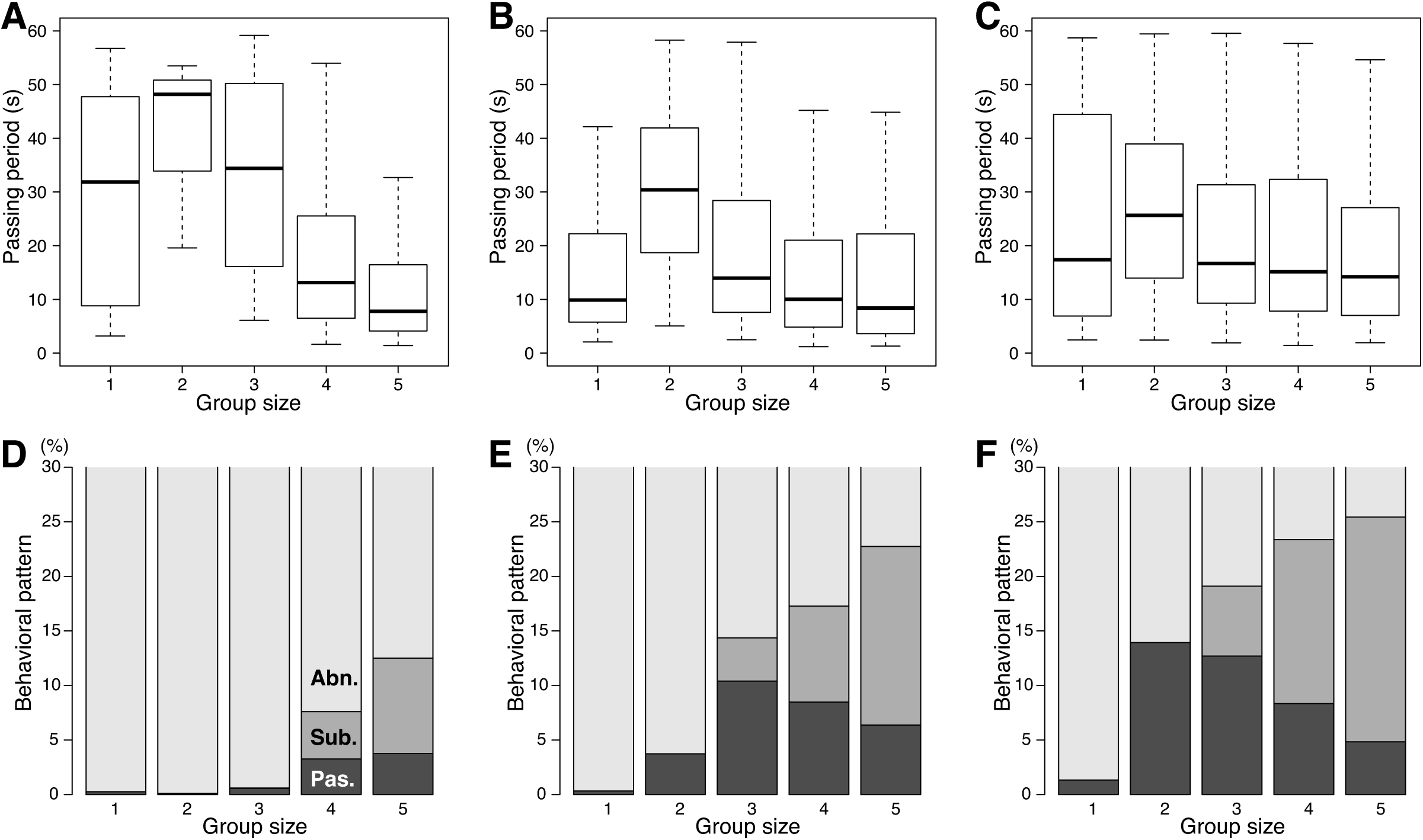
Simulation results for obstacle-passing period with a U-shaped obstacle. A–C: Obstacle-passing period depending on group size. D–F: Behavioral patterns after an ant group faces an obstacle. A and D: Small diversity in directivity(*κ* = 4.4). B, C, E, and F: Large diversity in directivity (*κ* = 0.9). Stochastic force generation as *p* = 0.25 was considered in C and F. Other parameters were set as *f* = 10, *γ* = 0.4, and γ_*t*_ = 7.5. Other notations and classification of behaviors are the same as those in Fig 3.

## Conclusion

Two types of colonies, those conditioned in an indoor artificial environment (I) and an outdoor natural environment (II), were analyzed for *F. japonica*. Their dependences for speed of group transportation on group size differed. Previously, this difference may have been ascribed to differences between species. In contrast, we concluded through mechanical model analysis that this difference in speed dependence can be explained by diversity in directivity for individual ants among colony members.

Additionally, results of numerical simulation suggested that broader diversity increased the success rate of obstacle navigation and reduced the obstacle-passing period by grouping, supporting navigation of more complicated terrain. Diversity in directivity was also observed in *Novomessor cockerelli* [19] and in *Solenopsis invicta* [26], where the trajectories of transportation by single ants were distributed *around ±*45*° −±*60*°*, although they were not discussed the effect of diversity in their reports.

It is interesting that there is a trade-off relationship in transportation speed and rate of success in obstacle navigation. The speed increased depending on group size in colony (I), which possessed less diversity in directivity. The model analysis suggested that this is unsuitable for obstacle navigation. If the ants’s habitat is in the open and allows for a long line of sight, there are fewer obstacles but the probability of being attacked by predators is higher. Thus, rushing into the nest is a good strategy. In contrast, the speed did not increase depending on group size in colony (II), which possessed broad diversity in directivity; this is suggested as being suitable for obstacle navigation in complicated terrain. In such environments, the probability of being attacked by predators is lower. Thus, the ants can give priority to reliable obstacle navigation rather than transportation speed in such cases.

Lastly, the simulation results revealed that stochastic force generation, whether generated coincidentally by friction with the ground or intrinsically by the system [11, 12], can also assist in navigating more complicated obstacles. The success rate could be further enhanced by using an informed-ant-attachment mechanism [3].

There are still open questions, such as how colony members learn the configuration around their nest and encode this information into diversity of directivity. Additionally, how ants whose preferential directions are opposite to the nest return to their nest remains unknown (these ants continued food transportation in a direction opposite to the nest over a distance of more than 1 m in the outdoor observations). Afurther question is whether distribution directivity depends on distance to the nest?

## Materials and Methods

### Ant preparation

Two types of colonies of Japanese wood ants, *F. japonica*, were prepared: ants kept in an indoor artificial environment, named colony (I), and ants that inhabited an outdoor natural environment, named colony (II).

Colony (I) was purchased from a vendor (Arinko Spot, Chiba, Japan), which collected a queen ant from sparse grassy ground located in Chiba prefecture, Japan, in July (nuptial flight season), 2015. Then the queen and pupae collected from another nest were kept in an artificial environment until the number of worker ants grew to a few hundred. The ants, including the queen and pupae, were transferred into an artificial nest made using a polystyrene chamber (176 × 80 × 32 mm), whose inner surface was plastered to maintain appropriate humidity and whose outer surface was covered with aluminum foil to maintain a dark environment. The nest was equipped with a single exit 7 mm in diameter. The nest was kept in another larger polystyrene*≥*chamber (300 × 200 × 50 mm) and was maintained under an environment of 26*°*C and 50 RH% humidity that was illuminated 12 h per day using LED light (800 lux). The ants were fed with sugar water or dissected mealworms every 3 days. The experiments were performed on the third day before feeding. One hour before the observations, the nest was placed into an arena made of polystyrene boards (1800 × 500 × 100 mm) as shown in Fig 2A. The observations were performed from July to October in 2016 in the indoor environment 27–28*°*C in temperature, 40–55RH% in humidity, and under conventional LED neutral white light bulb illumination (Luminous, LDAS40N-GM, Doshisha Corp.) by illuminating the observation stage.

Colony (II) was found in a plant pot (about 30 × 30 × 40 cm) located in a shaded place on a roof terrace on the sixth floor of building in Shinjuku-ku, Tokyo. Thus, this place was isolated from other colonies of *F. japonica*. Although it is known that *F.japonica* exhibits polydomy (i.e., a single ant colony forms multiple nest sites) [14],this colony exhibited monodomy (a single ant colony forming a single nest site), atleast during the observation season. The observations were performed in September 2013 in the outdoor environment 26–32*°*C in temperature, 40–55RH% in humidity, and in an unshaded place under fair or slightly cloudy weather.

The lengths of workers ranged over 4–6 mm in both colonies. Their living weight is known to be 5–6 mg [13].

### Food items

Food items consisting of Sakura-ebi (dried small shrimp *Sergia lucens*, around 40 mm in length and 120*±* 30 mg in weight) and Shirasu (boiled whitebait of sardine, around90*±*20mm in length and 90*±*20 mg in weight) were respectively prepared for colonies (I) and (II) to observe food item transportation. These were 5–10 times longer and 20times heavier than a worker ant.

### Observation of group transportation

Observation stages were placed at positions 150 cm away from the nest for colony (I) and 150–200 cm for colony (II), as shown in Fig 2A. First, a food item was placed in front of an ant foraging at a site on the observation stage. After the ant found the food item and recruitment was initiated, the food item baited by a single ant or bymultiple ants was transferred onto the observation stage. The observation stages were made of a polystyrene board (for colony (I)) or cardboard (colony (II)) and were equipped with a ‘rat-proof ‘structure to prevent additional ants from joining in the transportation of the target. Their behaviors were recorded using digital cameras(EXLIM EX-100, Casio Computer Co., Ltd. for colony (I); D-LUX4, Leica Camera AG for colony (II)) and saved as MPEG-4 videos or JPEG linear PCM photos.

### Obstacle

An obstacle made of polystyrene (W 6.8 × D 1.5 × H 3.7 cm) was prepared. Its lateral walls were coated with fluoropolymer resin (Fluon^®^PTFE, Asahi Glass Co., Ltd.) for colony (I), and with fragrance-free baby powder made of zinc oxide (baby powder,Pigeon corp., Japan) for colony (II) in order to prevent ants from climbing the walls.

### Data analysis

#### Images

Movies were converted into sequential TIFF format images at 10 fps in 8-bit gray-scale using image processing software (ImageJ [27]), and were then abstracted at the center position of the food item or the ants in each frame. The trajectories during transportation were obtained as demonstrated in Fig 4A.

#### Estimation of transportation speed

Speed of transportation was estimated as *d/T* using the period *T* that the food item moved a fixed distance *d* before the ants touched obstacles or the number of ants in the group changed (Figs 2B and C).

#### Estimation of obstacle-passing period

Periods for passing the obstacle were estimated as the difference between the time when one of the group ants touched a target area of the obstacle (an open circle denotes the touching point and the part shaded with oblique lines on the obstacle denotes the target area in Fig 2B) and the time when all group members transited across the line of the obstacle surface (denoted by a closed circle in Fig 2B).

#### Classification of obstacle navigation

Behavioral patterns when ant groups faced the obstacle were manually categorized into the following three types by watching the videos: the case where the ants succeeded in the task with the initial ant members and the initial arrangement(denoted by “Pas.” in Fig 3E), the case where at least one of the members left the food item and/or was substituted by another ant and the group succeeded in the task (denoted by “Sub.” in Fig 3E), and the case where all members abandoned the transportation task, releasing the food item (denoted by “Abn.” in Fig 3E).

#### Estimation of transportation direction

The angle of the transportation direction *θ*(*t*) at time *t* was defined using the direction of the position at time *t* + *τ* from the position at time *t* relative to the nest, as shown in Fig 2C. By moving the window of *τ* with the interval *δt* = 0.1 (equal to the image interval), a data set of *θ* values for a single trajectory was obtained. Then thedirectivity *θ*_0_ and the magnitude of fluctuations *σ* were estimated as explained in the following subsection. The value *τ* = 1.0 s was set so that *θ*_0_ could be captured more accurately and *σ* could be captured with moderate accuracy as follows. Various values of *τ* ranging from 0.1–2.0 s were tested as shown in S9 Fig. The mean value of the distribution of *θ* was saturated at around *τ* = 1.0 (S9 FigA), suggesting that larger values than this (denoted as (a) in S9 FigA) would be appropriate for capturing *θ*_0_ more accurately. If *τ* were too small, measurement errors would be detected (S9 FigB and C), leading simultaneously to worse estimation of *θ*_0_. On the contrary, if *τ* were too large, fluctuations in transportation directions would not be captured (S9 FigB and E). Thus, the range between them would be appropriate (denoted as (b) in S9 FigB). Therefore, *τ* = 1.0, which satisfied both conditions, was used.

Note that the magnitude of fluctuations would be underestimated in this scheme. It is, however, difficult to evaluate this magnitude because its dependence on *τ* exhibits a continuous curve as shown in S9 FigB. Therefore, we tested the effect of fluctuations with low resolution in terms of presence or absence. Without fluctuation, most trials failed in obstacle navigation. Thus, all simulations were performed with fluctuations.

#### Definition of directivity and fluctuation

From the data set *θ*(*t*) obtained by the above method, the distribution was analyzed as follows. The distribution was fitted with a normal distribution function using all the data first, e.g., as denoted by the blue dashed line in Fig 4B. Next, a normality test (Shapiro-Wilk test) was applied. If the *p* value was smaller than a level of significance (*α*=0.05 was used), rejection of normality was suggested and the data value farthest away from the mean value was removed (denoted as an outlier). Then the application of the normality test and outlier removal were applied until the distribution could fit the normal distribution (*p > α*, shown by the red solid line of Fig 4B). The data set of outliers corresponded with irregular movements such as sudden changes of the moving direction denoted by arrow heads and open blue circles in Fig 4A. Consequently, the final mean value was defined as the directivity *θ*_0_, and the standard deviations defined as the magnitude of the fluctuation *σ* were obtained.

#### Analysis of directivity

The directivity *θ*_0_ can be a measure of individuality of each ant, and was distributed around the nest direction for each colony as shown in Fig 4C and D. To characterize the distribution, mean values of *θ*_0_ and of the variance (*V*) were estimated using directional statistics [20] using the “circular” package in R [28]. The variance was defined in the domain of 0*≤V ≤*1. The value *V* = 0 represented all data pointing in the same direction, suggesting the smallest level of diversity. The value *V* = 1represented a uniform distribution in all directions, suggesting the largest level of diversity.

For application to the mathematical model in the discussion section, the distribution of *θ*_0_ was fitted with a von Mises distribution, a circular version of the normal distribution using the “circular” package in R [28]. The mean value, equal to that of the above circular statistics, and a concentration of 0*≤κ*, which is a measure of diversity, were analyzed.

### Numerical simulation

#### Derivation of rotation friction torque

In the Eqs (1) and (2), the speed *v*(*s*) of a position at a distance *s* from the center on the rotating rod with rotation speed *ω* is represented as *v*(*s*) = *sω*. Therefore, therotation friction torque was calculated by the following integration:

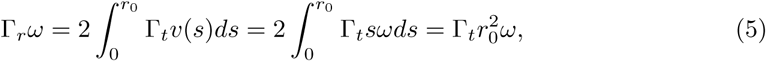

where 2*r*_0_ is the length of the rod.

#### Theoretical analysis of transportation speed using Eq4

Eq (4) and the order parameter *R*(*n*) defined by Eq (3) were statistically analyzed. The directivities for individual ants among a group were randomly generated according to a von Mises distribution with a mean parameter of 0 and a concentration parameter of *κ* =0, 4.4, and 0.9 for calculations corresponding to no diversity, low diversity, and high diversity, respectively, as shown in S6 Fig. Then 1,000 trials foreach group size were performed and distributions of *R*(*n*) and |**v**| were analyzed using their medians and box-and-whisker diagrams.

#### Simulation of transportation

Computer simulations using Eqs (1) and (2) were performed. For simplicity, inertia was neglected; namely, left-hand terms in Eqs (1) and (2) were approximated as 0. The Euler method was applied with a time step size of Δ*t* = 0.1 s. Initial positions of food items were set 3*r*_0_ apart from the center of the obstacle.

#### Simulation of obstacle navigation

A rectangular (Fig 5B) or U-shaped obstacle (Fig 5C) was placed in front of the transported food item so that the center of the food item faced near the center of the obstacle within the range of 2*r*_0_. In our observations, even when ants reached the wall of the obstacle, they persisted in moving in the same direction but failed to climb the wall owing to slipping; thus, the position did not move. Consequently, we assumed that the ants exerted a force from the obstacle equal to the normal vector according to Newton’s third law.

We judged the time when whole body of the food item passed through a line denoted by the dash-dotted line in Figs 5B and C, as the time when an ant group succeeded in navigation of an obstacle. The obstacle-passing period was then estimated using the same method as in the experiments. When the period exceeded 60 s, we judged that the group had abandoned the transportation task (‘Abn.’) in Figs 7 and 8. The critical period was estimated according to the experimental results, in which the period required for groups to succeed in the task was always less than 50 s. The category of ‘Sub.’ denotes the case where the initial direction of an ant’s first bite was unmatched with its preferential direction but where the group succeeded in passing the obstacle nevertheless. The category of ‘Pas.’ denotes the case where there was no directional mismatch and the group succeeded in the task.

## Supporting Information

**S1 Video. Tug-of-war in group transport.** Intervals between time stamps are 1/30 s. Time stamp 968, 998, and 1028 correspond to the photos of A, B, and C in Fig 1, respectively.

**S2 Video. Success in passing the obstacle.** Three ants in a group transported a food item and eventually passed the obstacle.

**S3 Video. Abandonment of obstacle navigation.** A single ant tried *to transport* a food item across the obstacle but abandoned the task, releasing the food item.

**S4 Video. Persistence of directivity.** A single ant was transporting a food item in a certain direction. Even when interrupted and moved to another location, the ant continued to transport the food item in almost the same direction.

**S5 Fig. Mismatch between the first bite direction and the ant ‘s preferential direction.** The ant pulled the food item in the body-backward direction first (red arrow) and then corrected the pulling direction to its preferential direction (blue arrow).

**S6 Fig. Order parameter** *R*(*n*) **depending on group size** *n*. (A) No diversity. (B)Low diversity. *κ* = 4.4. (C) High diversity. *κ* = 0.9. The values *R*(*n*) were calculated using Eq (3). The directivity values *θ*_0_ of individual ants were randomly generated from the von Mises distribution with the mean equal to 0 and with the concentration parameter *κ*. Thus, the values of *R*(*n*) among trials were distributed, and were obtained from 1,000 trials and represented by box-and-whisker diagrams.

**S7 Video. Simulation results for obstacle navigation with no diversity.**

The preferential directions of two ants were set to the direction of the nest (upper area in the movie). It took longer to finish obstacle navigation in this case. The directions of the traction force of individual ants (represented with magenta lines) fluctuated around their preferential directions (represented with black lines).

**S8 Video. Simulation results for obstacle navigation with high diversity (***κ* = 0.9). The preferential directions of two ants were set randomly relative to the direction to the nest. It was easy for the group ants with high diversity to succeed in the obstacle navigation task.

**S9 Fig. Estimation of transportation direction.** (A) Mean value of the distribution of the transportation directions *θ* depending on the time step *τ* used in calculations. (B) Standard deviation of the distribution of *θ* depending on *τ*. (C–E)Examples of transportation trajectories. The values *τ* =5, 10, and 20 were set for (C), (D), and (E), respectively. (a) An appropriate range for estimating *θ*_0_. (b) An appropriate rage for estimating *σ*.

## Acknowledgments

The authors wish to thank Professor H. Nishimori, Hiroshima University, and Mr. S.Hisamoto, Waseda University, for useful information on keeping ants and on the experimental setup.

